# MagnEspScope - DIY microscope with integrated magnetic system for magnetogenetic studies

**DOI:** 10.1101/2025.05.10.653253

**Authors:** A.S. Minin, M.A. Uimin, A.V. Belousova, I.V. Zubarev

## Abstract

Magnetogenetics is a promising field of biology devoted to the manipulation of biological objects by magnetic fields, where magnetic nanoparticles usually act as actuators. However, studying the response of biological objects is hampered by the fact that microscopes are usually made of ferromagnetic materials, making it difficult to work with them in powerful magnetic fields. This article focuses on building a DIY 3d printed microscope based on the ESPressoscope platform, integrated into a gradient magnetic system of permanent magnets. The article reveals the design, choice of materials and, using the example of cellular spheroids saturated with magnetic nanoparticles, demonstrates the application of MagnEspScope for the observation of magnetophoresis of biological objects.

## Introduction

There are many different ways to affect biological objects: light, mechanical action, electricity, ultrasound and, of course, countless chemical substances. Some of these methods are routinely used both in medicine and in basic research in biology, others are awaiting their introduction. The latter includes the application of magnetic fields. Of course, there are such methods of research as MRI and NMR, but in terms of exactly the impact on the body, in order to obtain some biological response, everything is much more complicated. Such a method as transcranial magnetic stimulation[1] is relatively widespread, which allows influencing the nervous system with powerful electromagnetic fields, but that is practically all.

Nevertheless, in recent years, new approaches have been appearing, conventionally united into one discipline called magnetogenetics [2]. The advantage of magnetic field is its ability to penetrate almost unlimitedly inside biological objects, which favourably differs from the method of optogenetics. However, optogenetics has become popular due to the fact that there is a large set of molecular mechanisms that can be affected by light, whereas no molecular targets for magnetogenetic effects have been found yet, except for some controversial experiments with ferritin [3] and similar molecules and the MagLov family of proteins[4], which has not yet found applications. Hence, the tools of magnetogenetics are various kinds of magnetic nanoparticles, which in one way or another are bound to various receptors and organelles in cells[5], after which they are affected by alternating or constant magnetic fields[6].

Among the complexities of magnetogenetics, one particularly complicates work at the cellular level: microscopy. All modern microscopes are made of ferromagnetic alloys, which are quite poorly compatible with strong gradient magnetic fields. Firstly, the microscope can be damaged, and secondly, a ferromagnetic object will distort the magnetic field. Accordingly, the microscope should be made of brass, aluminium or other materials, but such solutions on the market of scientific equipment are unreasonably expensive.

This paper focuses on the design and construction of a microscope from plastic parts printed on an FDM 3d printer. This microscope is based on the ESPressoscope [7] open-source project, refined for the specific task of magnetogenetics as applied to the effects of magnetic nanoparticles on cellular spheroids. For any scientist wishing to reproduce our results the protocols, schematics and 3d models of our developed design are available on the project’s GitHub page https://github.com/arteys/MagnEspScope.

## Materials and methods

### 3d printing

FDM 3d printing was carried out from PLA and PETG filament by Bestfilament (Russia) and NIT (Russia) on Ender 3v3SE printer, the used printing profiles are given in the auxiliary materials. PrusaSlicer version 2.9.2 was used to prepare 3d models for printing.

#### Assembly of the magnet system and microscope

To assemble the magnet system use NdFeB magnets, two sizes: 40×40×15 (magnetised on axis 15) and 40×20×10 (magnetised on axis 10). Sub-assemblies were assembled from the magnets: two large magnets and four small magnets into a printed case holding the magnets. Further assembly was carried out on an iron table to simplify the process. The assembly sequence was carried out in the order shown in the figure. Tools made of non-ferromagnetic materials like levers and spacer wedges were used during assembly. It should be remembered that permanent magnets of this size can generate a very significant mechanical force, posing a hazard to those handling them, so assembly should be carried out by qualified personnel.

**Figure.**
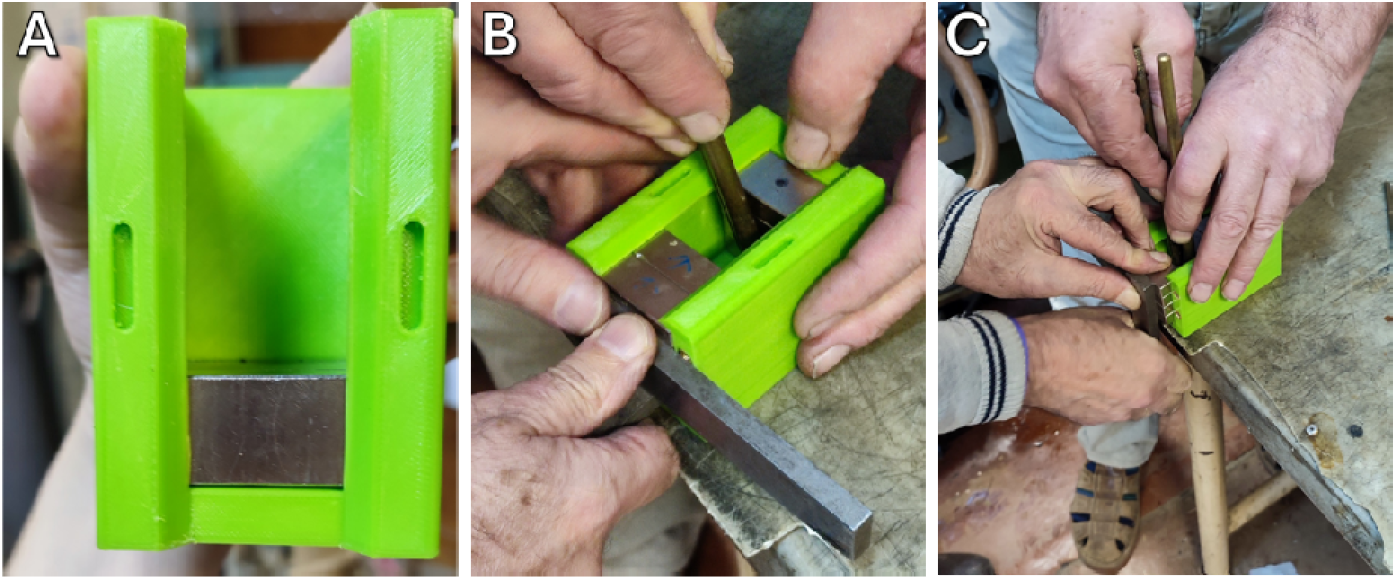
Stages of assembly of the magnetic system

The microscope itself, as mentioned above, is based on the ESPressoscope design, with a number of modifications. The 3d printing of the microscope was made from PLA plastic, and brass threaded inserts were soldered into the key holes, allowing for a more accurate and precise focus adjustment than the original design. Also the shape of the stage has been modified so that the microscope can be inserted into the magnet system The heart of the microscope is the ESP32-CAM controller with an OV2640 camera. The latest version of Matchboxscope software is installed on the controller. The illumination is a white LED mounted on top in a retrofitted illuminator. The LED is powered separately, not from the controller, as this leads to overheating of the latter. Communication with the controller was carried out via WiFi, also when the MagnEspScope is inside the cell incubator.

### Cell cultures and spheroids fabrication

The study was conducted on HEK-293t cells (human embryonic kidney epithelial cells) obtained from the Russian collection of cell cultures of the Institute of Cytology, Russian Academy of Sciences. The cells were maintained in DMEM nutrient medium (BioinnLabs, Russia) supplemented with 10% FBS (Biolot, Russia) in an incubator at 37°C and an atmosphere containing 5% carbon dioxide. Cells were tested for the absence of mycoplasma before the experiments.

Cell spheroids were made using the agarose microwell method using moulds from the MSLASPheroidStamp project [8]. For this purpose, a 2.75% agarose solution was melted and transferred into silicone moulds with a microwell size of 500 μm. After solidification, the moulds were transferred into the cells of a 6-well culture plate cell. Then, a suspension of HEK-293t cells in nutrient medium in the amount of 2000 cells per microwell was poured into them. Spheroids were formed within 24 hours.

To obtain spheroids from cells saturated with magnetic nanoparticles, cells were incubated with FeC-NH2 magnetic nanoparticles added to the nutrient medium at a concentration of 0.1 mg/ml before spheroids were made, after which spheroids were made in the manner described above.

Cell culture and cell spheroids were observed using an Olympus IX-71 inverted microscope.

### Image processing

Image processing was performed in the ImageJ (FIJI) package [9]. The images obtained with MagnEspScope were converted to 8-bit format, and the brightness and contrast were auto-tuned to make the details better distinguishable.

### Software

The magnetic system was modelled in FEMM version 4.2, which allows solving magnetostatics problems using the finite element method. The magnetic system and the microscope were modelled in freeCAD version 1.0.

## Results

### Magnetic system

The main idea of the MagnEspScope project is to build a microscope that allows us to observe the effects of mechanical forces acting from inside cells by means of magnetic nanoparticles. The force exerted on a nanoparticle by a magnetic field depends on the gradient of this field, not only on the strength. Accordingly, the microscope should be equipped with a magnetic system allowing to create a constant gradient magnetic field. Such a system can be made on the basis of an electromagnet, but permanent magnets allow to make it, in general, more powerful and at the same time energy-independent.

#### Magnet configuration

The purpose of creating this magnetic system is to study the effect of magnetic field on various biological objects, in the framework of this article on cell spheroids. These are usually cultured in culture plates and petri dishes of different formats. However, in order to recruit adequate statistics, it is necessary that the exposure to the entire experimental area is the same. Hence, petri dishes with a diameter of 28 mm were chosen as the most compact culture dishes used. Plate cells of course come in smaller diameters, but the plate itself is a rather bulky object.

To create the gradient magnetic system, the three permanent magnet blocks were configured in the manner shown in Figure A. These can be either solid magnets or sub-assemblies of individual magnets. This design allows for a uniform gradient at the mid-height of the system (Figure B).

**Figure.**
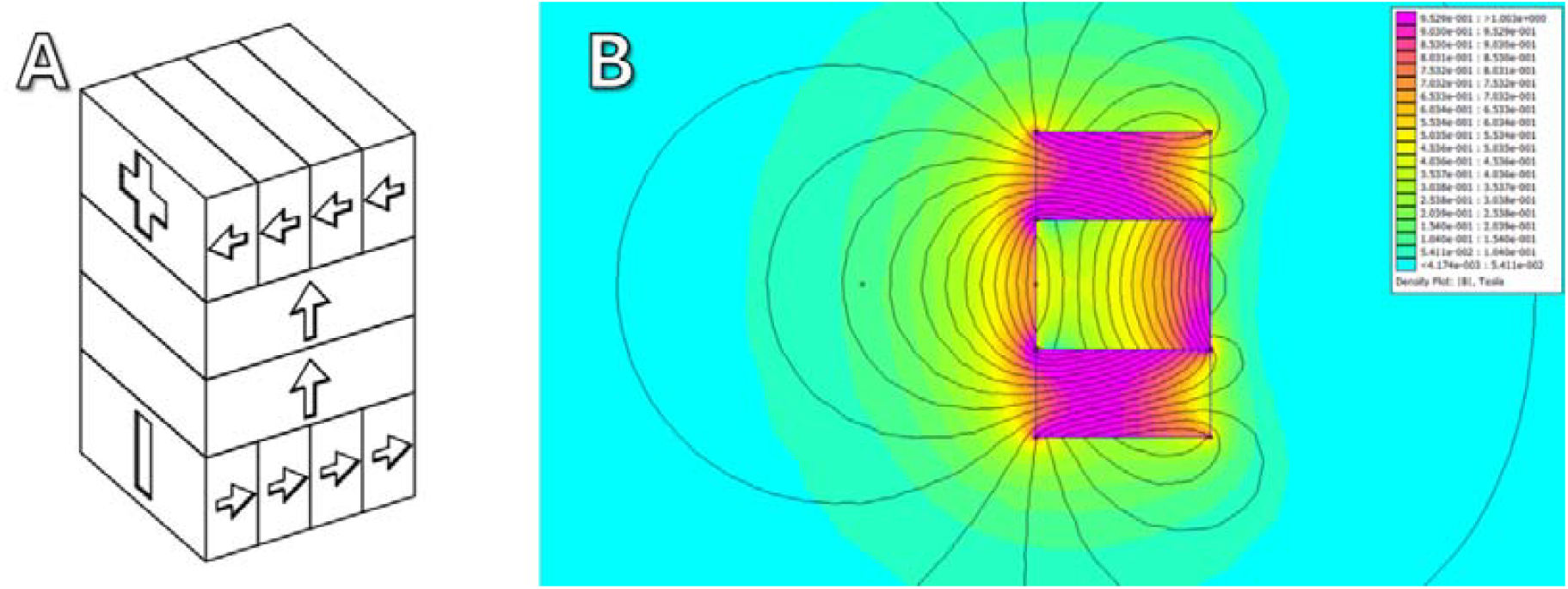
Schematic of the magnetic system design and magnetic field topology calculated in FEMM

#### Material selection

The magnets in a magnetic system tend to turn towards each other with opposite poles, resulting in a significant mechanical force. As magnetic systems of this kind are held as a unit by being glued in epoxy resin or by means of an external housing. The housing should be of non-ferromagnetic material: bronze, brass, aluminium alloys. However, to simplify prototyping and, in general, to make the design cheaper, we used a 3d printed enclosure. The most common material used in FDM 3d printing is PLA, but its notable disadvantage is creep [10], especially at elevated temperature. The first version of the magnetic system remained robust at room temperature, but began to deform when placed in a 37 degree cell incubator (Figure A). The second version was designed with a PETG housing, in addition, the housing was further reinforced, and together this eliminated the deformation problems (Figure B).

**Figure.**
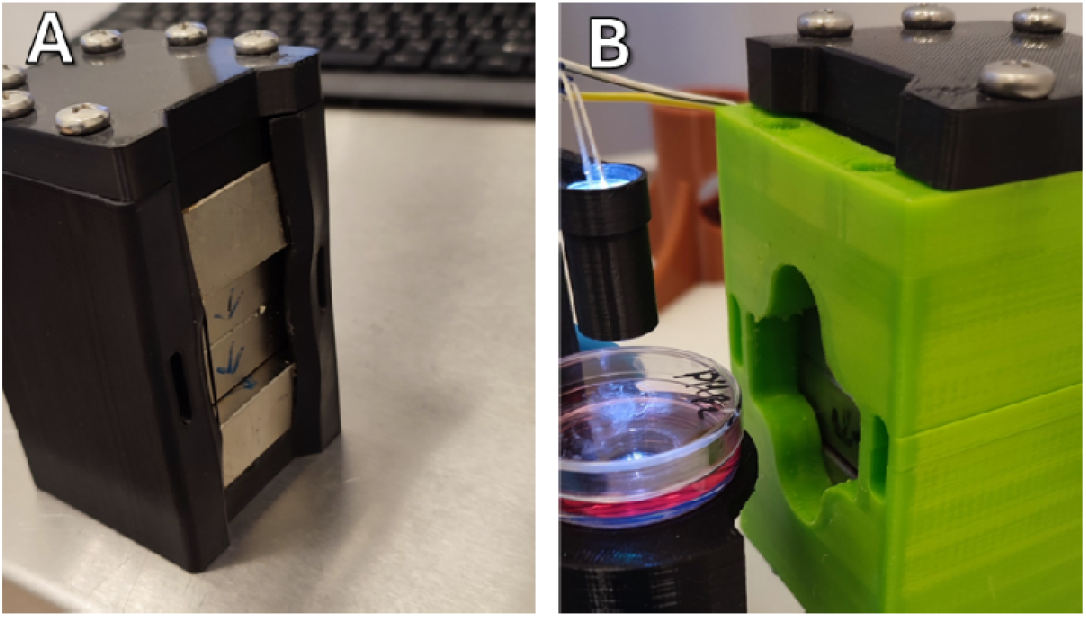
Deformed magnetic system with PLA body (A), after 24 hours in incubator at 37C and system with reinforced PETG body (B) periodically used for several months.

#### Magnetic field in the system

The magnetic field in the system was measured using a three-axis Hall sensor system and it was generally consistent with the original calculations. In the horizontal plane (A) of the petri dish the field ranged from 500 Oe to 2500 oE and the gradient was respectively 70 Oe/mm or 7 T/m (B), and vertically the range with a more or less homogeneous gradient was about 20 mm, which leaves quite a lot of room for manoeuvre (Figure C).

**Figure.**
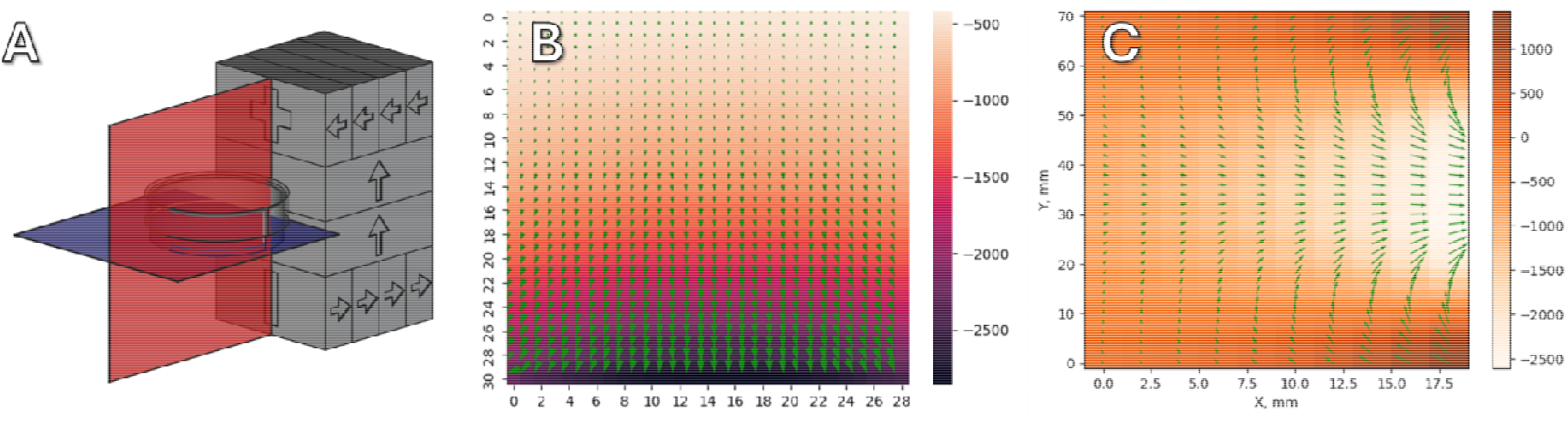
Results of field measurement in the gradient magnetic system. A - schematic illustration of the magnetic system. Blue denotes the horizontal plane, red - the vertical plane. Correspondingly B and C are the results of measuring the magnetic field topology in these planes.

**Figure.**
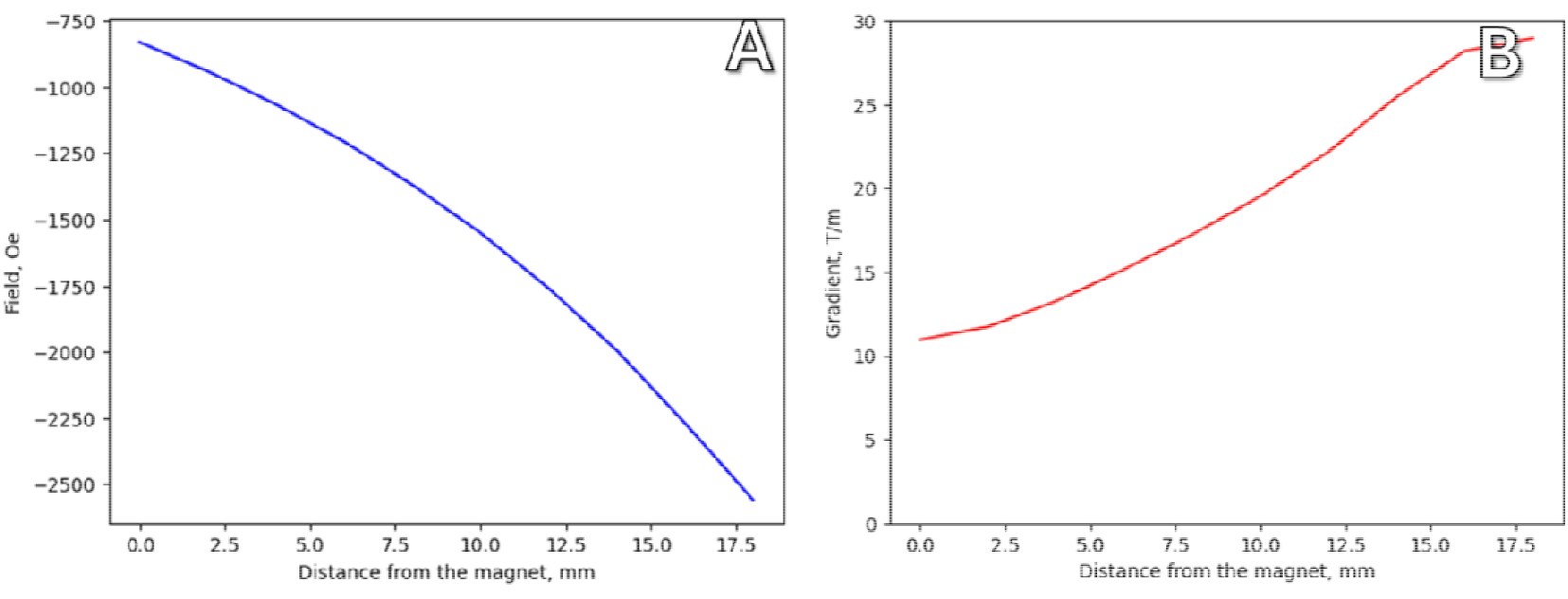
Magnetic field strength (A) and gradient (B) measured in the horizontal plane in the area of the bottom of the petri dish

### Microscope

Observation of objects in a strong magnetic field, especially with a noticeable field gradient imposes significant limitations on microscopy methods. The vast majority of standard microscopes are made of ferromagnetic materials, and their interaction with the field will, firstly, distort the field, and in the case of a large gradient may damage the delicate optomechanics of the optical system. It is of course possible to make the optical system from non-ferromagnetic metals, however, as in the case of the magnetic system housing, it does not seem economically rational in this case, hence the choice of FDM 3d printing as a technology.

There are a number of opensource DIY microscope projects where the basic structure is made of plastic and FDM printed. For example OpenFlexture, which utilises the elasticity of plastic to move the specimen in the XY plane [11] or PUMA, an advanced microscope with an integrated AR display [12]. Both of these projects are brilliant examples of the use of FDM technology, but their designs are rather bulky. However, there is another microscope that is as compact as possible, albeit primitive, called ESPressoscope, its name plays on a play on words between the size of an espresso cup and the underlying ESP family controller[7].

First of all, we slightly modified the slide of the basic version of the microscope to fit a petri dish on it (Figure A) and compared the image obtained with the ESPressoscope and a conventional inverted microscope (Figure B and C)

**Figure.**
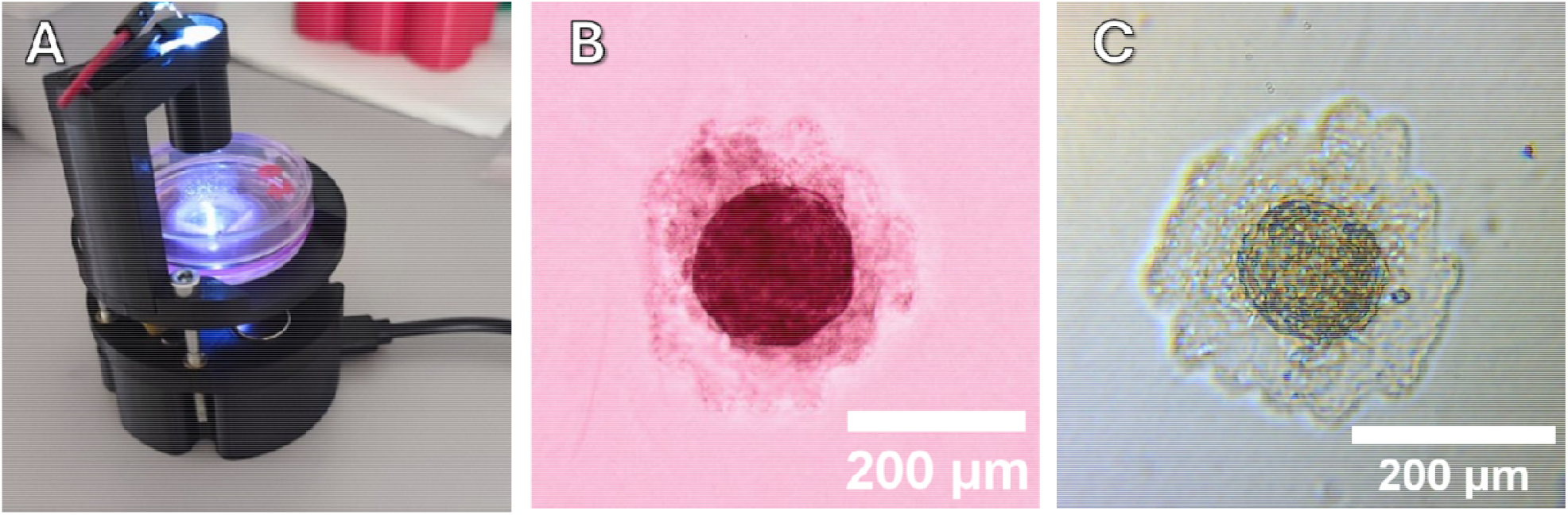
Basic version of ESPressoscope (A), comparison of images of cell spheroids from HEK-293t in agarose microwells obtained in ESPressoscope and in conventional microscope

The image obtained with the DIY microscope was, of course, significantly inferior in quality and detail to the conventional microscope, but the outlines and general details of the spheroid are quite distinguishable. Next, the ESPressoscope was modified, its shape was suitably changed, which allowed it to be mounted as close to the magnetic system as possible (Figure A and B).

**Figure.**
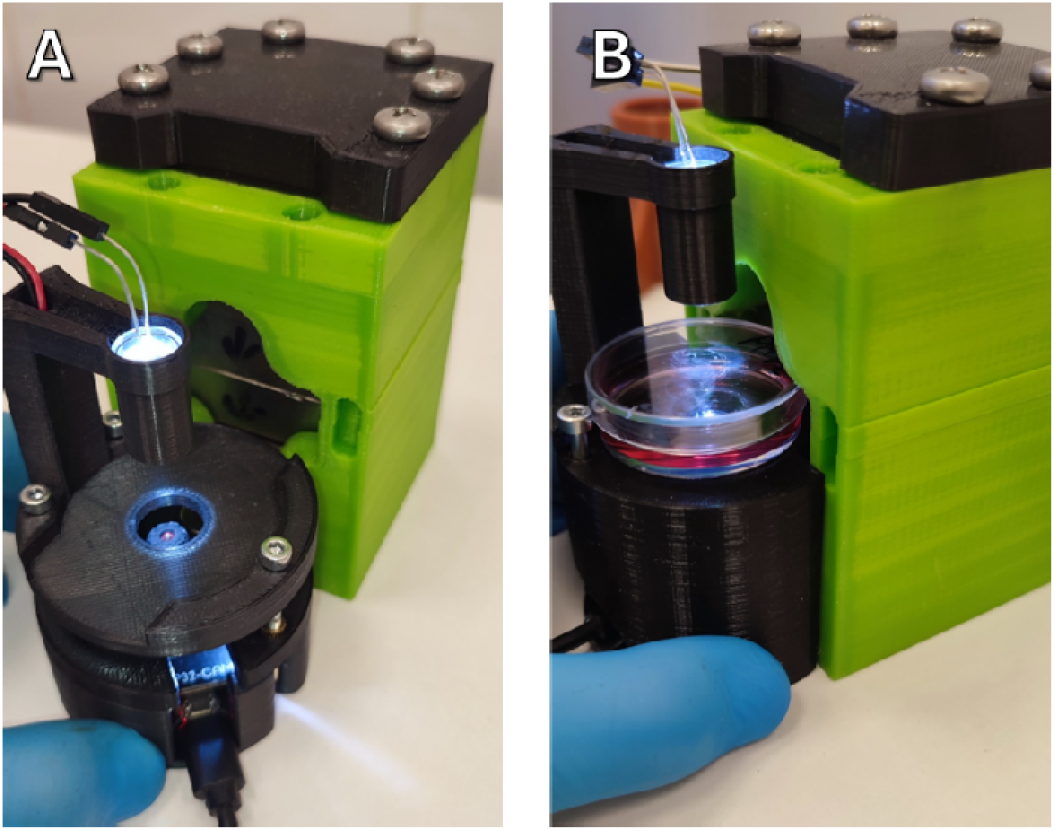
Basic version of the microscope and modified for installation in a magnetic system

### Magnetophoretic motion of spheroids

Finally, a suspension of cell spheroids from HEK-293t culture saturated with magnetic nanoparticles was placed in a petri dish mounted in the MagnEspScope. The main disadvantage of a microscope of this design is the lack of accurate positioning in the XY plane. While the focus (Z plane) can be adjusted quite easily using screws, movement in the XY plane is manual and requires extremely careful movements.

Nevertheless, it is possible and a cellular spheroid with an interesting shape was caught in the field of view, after which the microscope was carefully slid into the magnetic system. Then frames were taken every 10 seconds to capture the spheroid’s movement over time (Figure).

**Figure.**
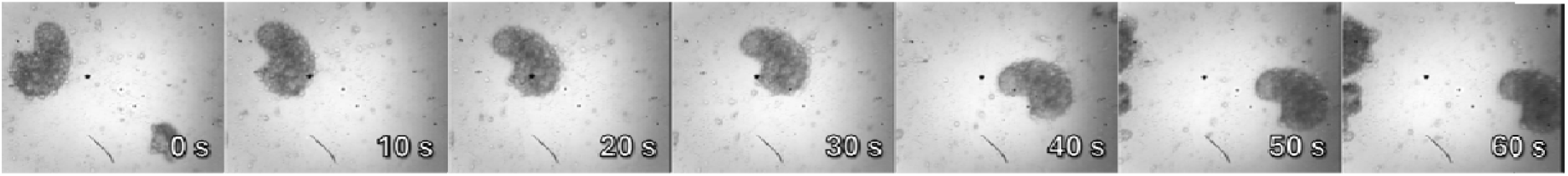
Movement of a spheroid from HEK-293t cells saturated with magnetic FeC-NH2 nanoparticles in a magnetic field (the magnet is located on the right), recorded using MagnEspScope.

The images obtained clearly allow us to judge the process of magnetophoretic movement of cell spheroids in a magnetic field.

## Conclusion

We have designed and constructed a microscope containing a minimum number of ferromagnetic parts, integrated into a gradient magnetic system (strength up to 2500 kOe with a gradient of 7 T/m). This microscope allows solving problems of magnetogenetics, observing the effect of magnetic fields on biological objects saturated with magnetic nanoparticles.

The design of the microscope is simple enough to be printed on a hobby-grade FDM 3d printer, and all drawings and protocols are available on the project’s GitHub page, allowing access for other scientists to reproduce our results.

## Acknowledgements

The reported study was funded by the Russian Science Foundation Grant #22-74-10041. The work and characterization of Fe@C nanoparticles was performed within the framework of the state assignment of the Ministry of Education and Science of Russia for IFM UB RAS

